# Recurrent neural networks as neuro-computational models of human speech recognition

**DOI:** 10.1101/2024.02.20.580731

**Authors:** Christian Brodbeck, Thomas Hannagan, James S. Magnuson

## Abstract

Human speech recognition transforms a continuous acoustic signal into categorical linguistic units, by aggregating information that is distributed in time. It has been suggested that this kind of information processing may be understood through the computations of a Recurrent Neural Network (RNN) that receives input frame by frame, linearly in time, but builds an incremental representation of this input through a continually evolving internal state. While RNNs can simulate several key *behavioral* observations about human speech and language processing, it is unknown whether RNNs also develop computational dynamics that resemble human *neural speech processing*. Here we show that the internal dynamics of long short-term memory (LSTM) RNNs, trained to recognize speech from auditory spectrograms, predict human neural population responses to the same stimuli, beyond predictions from auditory features. Variations in the RNN architecture motivated by cognitive principles further improve this predictive power. Moreover, different components of hierarchical RNNs predict separable components of brain responses to speech in an anatomically structured manner, suggesting that RNNs reproduce a hierarchy of speech recognition in the brain. Our results suggest that RNNs provide plausible computational models of the cortical processes supporting human speech recognition.

## Introduction

The computations underlying early auditory representations can be approximated as convolutional filters. Such mechanisms are typically studied using spectro-temporal receptive fields (STRFs) (Aertsen et al., 1980). A majority of midbrain and early cortical neurons can be well described with such models, especially when incorporating some specific nonlinearities (Escabí and Schreiner, 2002; Fishbach et al., 2001; Singer et al., 2018; Williamson et al., 2016). However, such mechanisms seem inadequate for capturing linguistic representations such as words. For example, STRFs have fixed temporal integration windows. In contrast, there is no hard upper limit to how long words can be, and words can be pronounced and recognized at drastically different speech rates.

Recurrent neural networks (RNNs) successfully solve such pattern detection problems in sequences that require integrating information over time (Elman, 1990), and may thus provide promising models for the neural mechanisms for human speech processing (Graves et al., 2004; Yi et al., 2019). RNNs are highly incremental: they fully integrate the current input into their internal state at each processing step. This may make them particularly suited for modeling human speech perception, which is highly incremental (Allopenna et al., 1998; Brodbeck et al., 2022; Christiansen and Chater, 2016). Finally, long-short-term memory (LSTM), a particularly successful RNN architecture (Hochreiter and Schmidhuber, 1997), may resemble the architecture of cortical microcircuits (Costa et al., 2017).

RNNs trained on phoneme symbol sequence inputs can simulate human sensitivity to phonotactic regularities in speech (Cairns et al., 1997; Donhauser and Baillet, 2020). RNNs developed for automatic speech recognition can process real speech acoustic inputs (Graves, 2012; Graves et al., 2013; Graves and Jaitly, 2014). However, these RNNs are not adequate models of human speech perception for several reasons: (1) these models typically process speech forward and backwards (with the signal available in an image-like fashion), whereas humans are presumably limited to forward processing by the natural progression of time; (2) the RNNs’ task is typically to transcribe a spoken input to a grapheme sequence, which is subsequently combined with an extraneous language model for transcription to words, whereas humans are thought to recognize phonological forms and words directly from speech, with simultaneous constraint from linguistic knowledge.

Recently, the RNN architecture has been adapted as a more realistic model of *human* speech recognition (Magnuson et al., 2020). An RNN was trained to recognize words, represented in the output as semantic feature vectors, directly from acoustic spectrograms. A simple architecture, consisting of a single hidden layer of LSTM nodes, was deliberately chosen to improve the potential for explainability. This model exhibited several characteristics of human speech recognition, such as the fine-grained time course of competition between words with overlapping phonology (Allopenna et al., 1998), and its hidden units exhibited phonetically organized responses to speech similar to those observed in human cortex (Chang et al., 2010; Mesgarani et al., 2014), despite not being trained on phonetic targets.

Here we ask whether the temporal dynamics of computations in such an RNN also come to resemble the time course of human neural responses to speech. We address this by developing RNNs that can recognize words from the same acoustic stimulus sequences heard by participants in a magnetoencephalography (MEG) experiment, and predicting MEG responses from the RNNs’ hidden unit activity (Figure 1). We compare the effects of different architectural decisions on the predictive power for brain activity to develop an RNN that most resembles human speech recognition.

**Figure 1.**
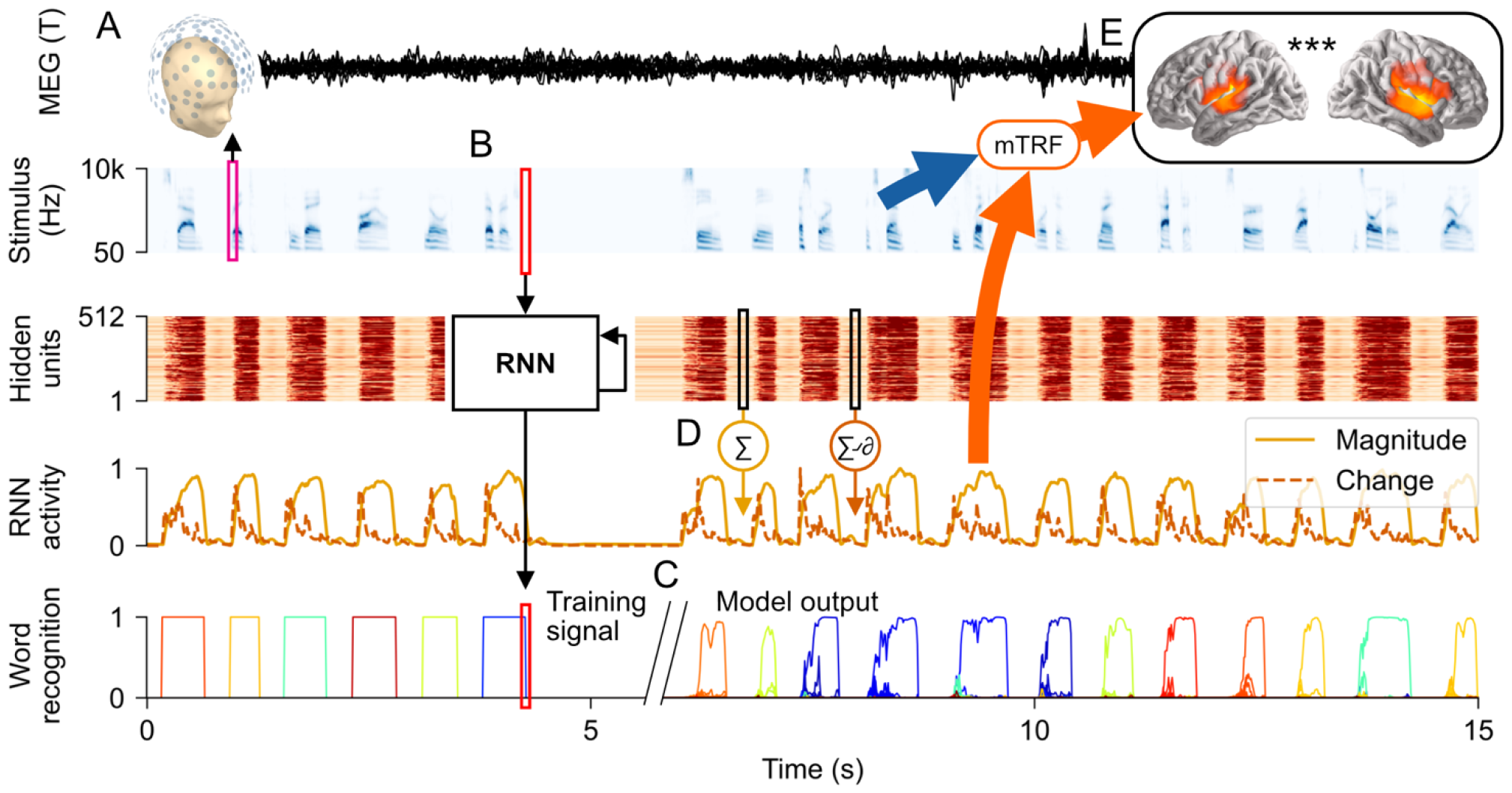
General design: predicting brain activity from a recurrent neural network (RNN). (A) Human participants listened to spoken words, while magnetic fields were measured with magnetoencephalography (MEG). (B) A recurrent neural network (RNN), trained to recognize words in arbitrary sequences of words and silence, processed the same stimulus sequence that each human participant heard. (C) The RNN was trained to output the word that is currently being heard, for the whole duration of the word (“Training signal”; each color represents a different word in the output space). In practice, this is impossible in the early time course, because information about word identity is distributed over time in the acoustic input. Instead, the RNN tends to activate several possible candidates before settling on the right word (“Model output”; words are sorted phonetically, thus words with similar color have a similar onset). (D) RNN activity over time was quantified as the sum of hidden unit magnitude and the sum of activity increases. (E) These two signals were then used to predict the source-localized MEG responses from each participant, while controlling for the predictive power of acoustic features (a gammatone spectrogram, an acoustic onset spectrogram, and word onsets). Brain responses were predicted through multivariate temporal response function (mTRF) models.

In recent years it has been reported that the most proficient language models are also the best predictors of human brain activations (Schrimpf et al., 2021). Because Transformers (Vaswani et al., 2017) currently dominate other models on all language tasks, including speech transcription (Gulati et al., 2020; Karita et al., 2019; Srivastav et al., 2023), it would be tempting to infer that they are inherently more “brain-like” (Li et al., 2023). However, this is liable to the confound that Transformers are more computation-efficient on current silicone hardware, which enables deeper and wider models to be trained on more data. To our knowledge, there has been no systematic comparative study of Recurrent and Transformer architectures as predictors of human speech activations, that controls for the number of floating point operations during training. Furthermore, the attention mechanism, which makes transformers so efficient, inherently entails the ability to compare information from all time steps in the input in parallel. Transformers thus circumvent a critical challenge that the human brain must solve: Transformers effectively process sound sequences as static images, whereas the auditory nerve provides auditory input to the brain strictly in the order in which it occurs in time. Our work thus demonstrates an alternative to predicting brain activity from state-of-the-art artificial intelligence engineering models (Caucheteux et al., 2023; Li et al., 2023). Instead, we purposefully use artificial intelligence tools to design simple model architectures to relate to human cognition.

## Results

To simulate human speech recognition, we trained LSTMs to recognize words from a gammatone spectrogram, a representation of sound that mimics the human peripheral auditory system. The input was always a continuous spectrogram, constructed by concatenating words and periods of silence. The trained output signal was a zero vector during silence, and a word-specific vector (differing depending on the output space, see below) for the entire duration of each word. All models were trained to recognize 2985 words, selected to approximately include the lexical neighborhoods of the 1000 words which served as stimuli in the MEG experiment. For model training, each word was spoken by 15 synthetic talkers in addition to the human talker used for the MEG experiment. For each talker, a different 1/16^th^ of words were withheld during training to serve as targets for evaluating model performance (i.e., the 1/16^th^ withheld during training with a specific talker were not withheld with the other 15 talkers; note that it is not possible to test the model on completely novel words because the mapping from speech to semantics is mainly arbitrary).

### Models with sparse output space predict human brain responses better than models with pre-structured semantic output space

What is the format of the target representation in human word recognition? One conjecture is that word recognition directly maps acoustic patterns to a semantic space (Gaskell and Marslen-Wilson, 1997). Furthermore, lexical co-occurrence based vector spaces have been described as candidates for psychologically realistic representations of lexical semantics (Lund and Burgess, 1996; Turney and Pantel, 2010). A model implementing this architecture is shown in Figure 2-A (left side): a dense layer projects the RNN output into the GloVe vector space (Pennington et al., 2014), and the network is trained using the mean squared error (MSE) loss function. Alternatively, sound may be linked to an abstract intermediate level of *wordforms* before being linked to semantics (Kapnoula et al., 2024). This was implemented using a localist (one-hot) output space, in which the output vector has the size of the lexicon, and each word corresponds to one of the elements of this vector. As is common for classification problems, the RNN was connected to the output through a dense layer with a sigmoid activation function, and was trained using binary cross-entropy (Figure 2-A, right side). An additional possibility is that the meaning of each word corresponds to a sparse set of features; this was implemented using Sparse Random Vector (SRV) output (Magnuson et al., 2020), similar to the localist output except that each word corresponded not to a single vector element, but to 10 randomly selected elements of a vector of length 300 or 900 (the original model used 10 out of 300 for a 1000 word lexicon, thus 10 out of 900 provides the same level of sparsity as the original).

**Figure 2.**
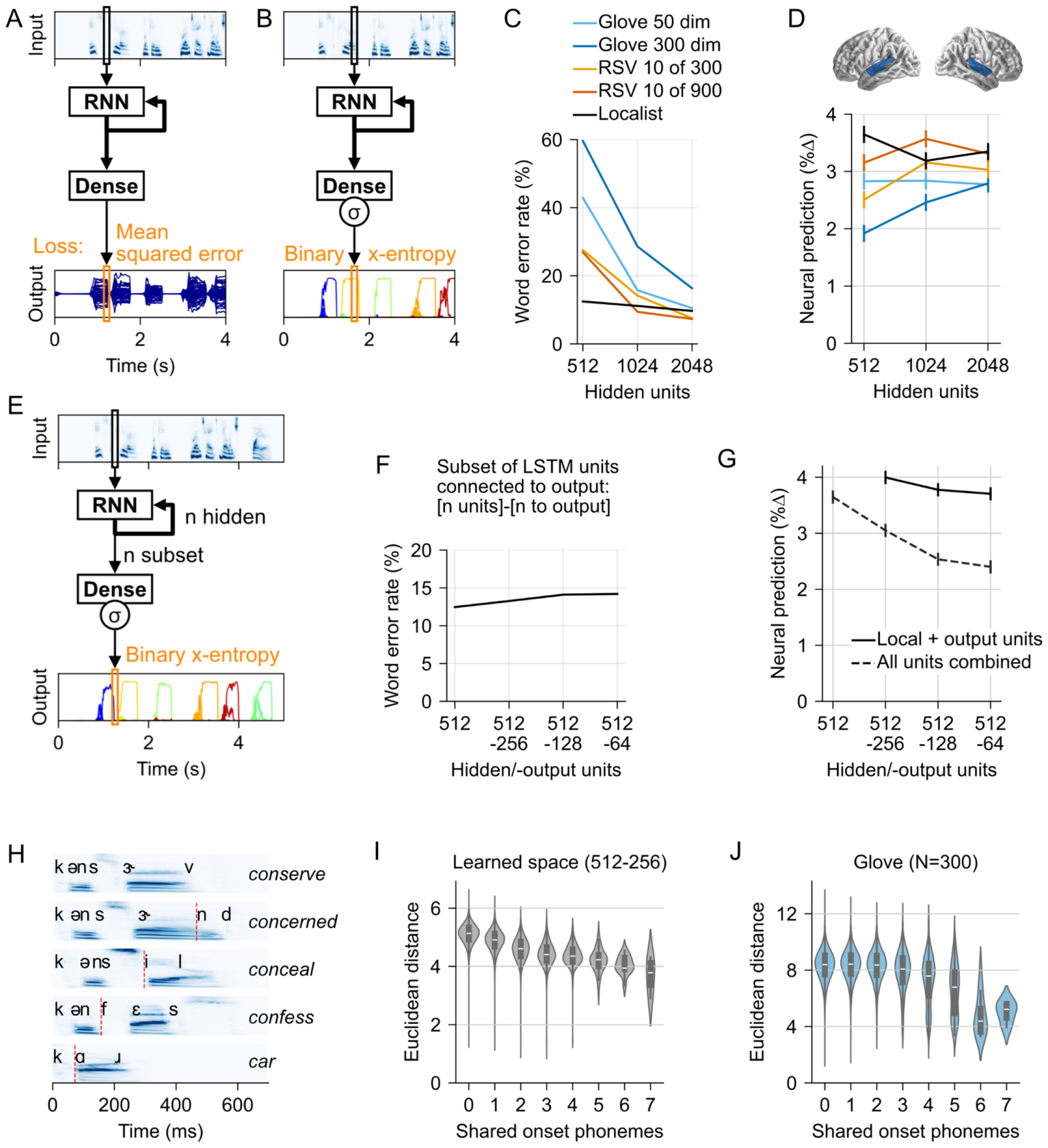
Models for sparse output spaces better predict neural responses, and learn phonetic structure of the input lexicon. (A) Architecture for RNNs with semantic output space. Word targets are dense vectors in a semantic vector space (GloVe). Each word has non-zero values in most or all output elements (blue lines). Output is evaluated using mean squared error (MSE). (B) RNN with sparse output, where targets are binary vectors, and each word is defined as 1 element (localist; shown) or 10 elements (sparse random vectors). In the illustration using a localist output space, each word leads to activation dominated by a single output element (different elements are distinguished by color). (C) Word error rate as a function of the numbers of hidden units, showing that all models can learn the task given enough hidden units. (D) Predictive power of each model for brain activity (quantified as % improvement in explained variance over an acoustic baseline model). All RNN models significantly add predictive power to the auditory-only model, but sparser RNN models have consistently higher predictive power than semantic vector space models. Error bars represent the within-subject standard error of the mean (Loftus and Masson, 1994). (E) Modification of the localist model to reduce the number of trained parameters. Only a subset (*n subset*) of all RNN nodes (*n hidden*) is connected to the output layer, leading to a substantial reduction in the number of weights in the fully connected layer between RNN and output space. (F) Word error rate as a function of how many hidden units are connected to a localist output. (G) The subsets of units that are or are not connected to the output explain complementary aspects of brain signals: Predictive power for brain activity decreases when all hidden units are combined to create a single predictor (dotted line). However, predictive power increases when separate predictors are created for representing the “local” hidden units that are purely recurrent, and the “output” hidden units that are also connected to the output layer. (H) Words commonly share the same acoustic-phonetic beginnings and can only be identified by considering all information across time. The plot illustrates this for *conserve* (first line): words lower in the graph share fewer word-initial phonemes. The point at which each word starts to differ from *conserve* is marked with a red line. (I) This acoustic-phonetic structure is reflected in the learned output mappings of localist models (dense layers): Words that share more onset phonemes are also located closer together in the output mappings. The graph shows the pairwise distance between words (x-axis) as a function of how many onset phonemes they share (y-axis). (J) Such acoustic-phonetic structure is not found in the GloVe output space.

When given a large enough number of hidden units, RNNs were able to learn each of the output representations (Figure 2-C), reaching ≤ 16% word error rate with 2048 hidden units. At lower numbers of hidden units, the denser output models performed notably worse.

Network performance did not strongly correspond to how well activity in the different RNNs predicted human MEG responses to speech. Figure 2-D shows the predictive power of each model for brain activity localized to the superior temporal gyrus of both hemispheres. Predictive power was quantified as the improvement in explained variance from adding the RNN activity predictors to an auditory baseline model (Brodbeck et al., 2020), i.e., 0% corresponds to the predictive power of the auditory model alone, and each data point in Figure 2-D corresponds to the predictive power of a model combining the auditory model with the hidden unit activity from an RNN model. Note that while improvements may seem small, this is variance *uniquely* attributed to the RNN, and does not include variance that is shared between RNN and auditory predictors. Importantly, the improvements are highly reliable across subjects, as indicated by the error bars.

The models’ predictive power for brain data varied by the number of hidden units and target space (*n-hidden* × *target-space*: *F*(8,269)=10.55, *p*<.001). Overall, the pattern of results suggests that the main feature that makes a model a better predictor of brain activity is the sparsity of the target space, with sparser target spaces performing better than the denser spaces. The densest target was GloVe, in which each dimension varies for all words; SRVs are less dense, with only 10 non-zero values per word, while localist targets have exactly 1 non-zero value. Critically, sparser models always outperformed the semantic vectors. The localist model was numerically better than the GloVe models at all levels, although this difference was not significant with GloVe-50 at 1024 units (Localist vs. Glove-50: 512: *t*(17)=3.63, *p*=.002; 1024: *t*(17)=1.84, *p*=.084; 2048: *t*(17)=2.82, *p*=.012; Localist vs. Glove-300: 512: *t*(17)=6.49, *p*<.001; 1024: *t*(17)=3.36, *p*=.004; 2048: *t*(17)=2.93, *p*=.009). However, at 1024 units, the 10 of 900 RSV model outperformed the GloVe models (Glove-50: *t*(17)=4.37, *p*<.001; Glove-300: *t*(17)=6.41, *p*<.001).

### Separating local and output units increases predictive power

The number of trainable parameters in the models strongly depends on the number of hidden units. Overall, the number of trainable parameters did not correspond strongly to the models’ predictive power for brain data (Figure 2-D). Nevertheless, at each level, the localist model had a disproportionately larger number of trainable parameters due to the dense mapping from hidden units to 2985 output dimensions (512→2985: 1,528,320), compared to the other models (e.g., 512→300: 153,600). The number of free parameters was reduced by connecting only a subset of RNN hidden units to the target vector (Figure 2-E). For example, a model designated 512-256 had 512 hidden units, of which only the first 256 were connected via dense mapping to the localist target vector (256→2985: 764,160 trainable parameters), while the remaining 256 hidden units were purely local, i.e., they served as recurrent units but could not directly affect the network output. These models with drastically reduced number of parameters only saw a small decrease in word recognition performance (Figure 2-F).

This division of units into different families (output vs. local) allowed an extension of the model for neural data, as the temporal dynamics in local and output units could be entered as separate predictors. This led to a significant improvement of predictions (Figure 2-G, solid lines; 512-256 vs. 512: *t(17)*=4.83, *p*<.001). In contrast, if local and output nodes were treated as a single set of units for creating a single predictor, predictive power decreased (Figure 2-G, dashed line). This suggests that the two types of nodes develop temporal dynamics that correspond to separable components in human neural responses. Whereas temporal dynamics of “output” units are directly constrained by the shape of the training signal, “local” units may develop task-appropriate activation patterns more freely (for anatomically separable components see *The layers of deep models predict flow of activity from auditory to higher level brain areas* below).

### Localist models learn acoustic-phonetic space

The semantic GloVe space serves as a low dimensional embedding and thus imposes a pre-defined structure on the lexicon. In localist models, the dense mapping from RNN hidden units to the 2985 word vector can similarly be interpreted as an embedding, as it encodes the 2985 words in a lower dimensional space (the RNN output units). Thus, one reason for why the localist models develop more human-like temporal dynamics may be that they learn a lexical neighborhood structure that is more reflective of human perception.

During word recognition, humans initially activate multiple lexical candidates (Allopenna et al., 1998). These early activations are determined by phonological similarity, as the first few phonemes of a word are usually compatible with multiple words. For example, upon hearing /bi/, listeners may consider both a beetle /bitƏl/ and a beaker /bik□/. Figure 2-H illustrates this for “conserve” (top), with competitor words that share different numbers of onset phonemes (the red line indicates the point at which they start to differ from “conserve” based on a phonological segmentation). This co-activation of words is driven by the acoustic-phonetic overlap, and is independent of any semantic relationship between the words. However, a semantic space like GloVe constrains such co-activation by its internal structure (Gaskell and Marslen-Wilson, 1997). For example, if *beaker* and *beetle* were on opposite sides of the space, they could not truly be both considered. The best the model could do would be an output corresponding to a point somewhere between the two words (which may incidentally correspond to a third, phonologically unrelated word).

We hypothesized that localist models may develop more human-like temporal dynamics by learning a lexical space that is structured such that it allows human-like co-activation of phonetic competitors. In such a space, words that are phonetic competitors ought to be closer to each other than words that are not. This was indeed borne out by the columns of the learned dense layer mapping RNN output to lexical identity. Figure 2-I shows the average pairwise distance between words in this space for the 512-256 unit model, as a function of acoustic-phonetic overlap, quantified as the number of shared onset phonemes. The correlation between pairwise distance and phonological overlap was highly significant (given the large number of word pairs: *r*=−.19, *p*<.001) and was strong even among words with 4 or fewer shared onset phonemes (*r*=−.19, *p*<.001). Figure 2-J shows the same analysis of the GloVe space for comparison. Here, the correlation is very low for 4 or fewer shared onset phonemes (*r*=.00) and only increases for words which share 5 or more onset phonemes (*r*=−.26). This latter increase is likely because words that share the same root are at the same time semantically related and phonological competitors (e.g., story/stories; breath/breathy; complain/complaint; vision/visual, truck/trucker, …).

### Deep models are more human-like

Hierarchical depth was introduced by stacking multiple RNNs, i.e. the output of the first RNN is the input for the second RNN and so forth (Figure 3-A). The most human-like shallow model served as a baseline (512-256, with 1,948,841 trained parameters). The number of hidden units for deep models was determined by selecting the multiple of 64 that led to the number of trainable parameters closest to 2,000,000. GloVe models were retained in this analysis because the addition of depth may allow more human-like acoustic-phonetic competition in lower levels, which are less directly constrained by the output space. Deep GloVe models might thus be better predictors of human brain activity by developing phonetic competition in lower layers, and semantic activation in higher layers.

**Figure 3.**
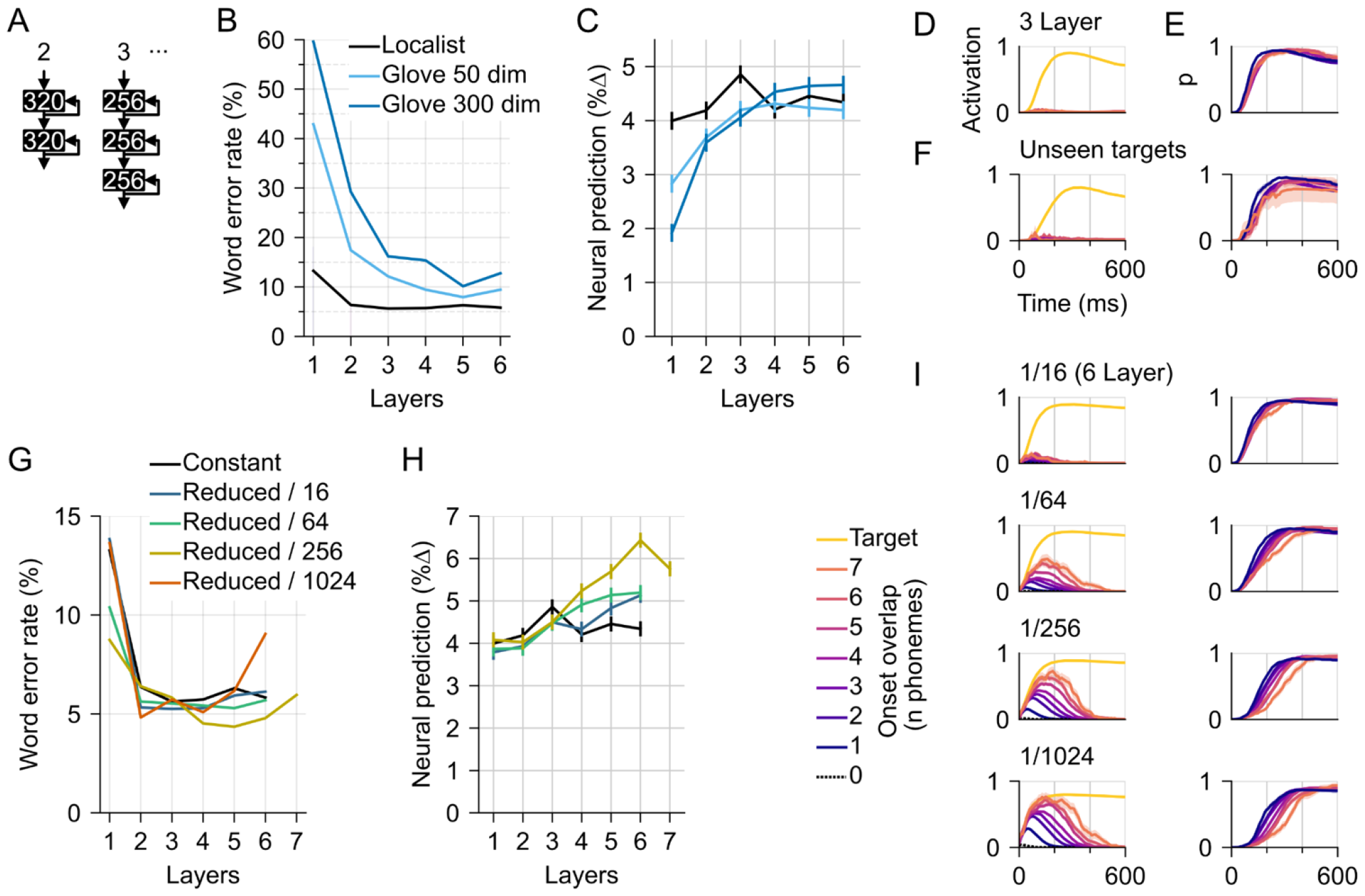
Depth and phonetic competition improve performance and neural prediction. (A) depth was added by stacking RNNs while controlling the number of trainable parameters (the number of hidden units per layer is indicated in white). (B) Deeper models have lower word error rate. (C) Deeper models also have better predictive power for brain responses. (D) Activation of cohort competitors is limited (shown for the best-performing localist model with 3 layers). The plot shows model output values in the localist space as a function of time since word onset. Shown is the output for the target (yellow), and the average for other words, with color indicating how many word-initial phonemes they share with the target. Only data for the human speaker tokens are shown. (E) The relative activation of the correct target, i.e., activation of the target divided by the sum of the total activation in the output space, at each time point. Color indicates the longest competitor, i.e., how many word-initial phonemes the target shares with at least one other word in the lexicon. Words that share more phonemes with competitors should take longer to uniquely recognize. Here, the correct target dominates from early on, regardless of the competitor environment. (F) Same as D and E, but for a subset of 1000 target tokens that were never presented during training (in a separately trained model; data for the same tokens also used in the MEG experiment). While target activation is delayed compared to D, the activation of competitors with shared word onsets is similarly low. This suggests that overlearning of specific tokens is not the cause for the absence of competitor activation. (G) The loss function was modified to reduce the penalty incurred by activating non-target words early during word presentation. The modified loss function led to improved WERs, and (H) substantially increased predictive power of deep models. A 7 layer model was tested for the *c*=256 reduced loss to determine whether the rising trajectory would continue. (I) The modified loss function also led to more human-like activation of cohort competitors early during words (left column), and delayed the point at which activation of the target exceeded other words, with the expected gradation by how many onset phonemes competitors share with the target (right column), more consistent with acoustic ambiguity of the word onsets (data from 6 layer models).

In terms of word recognition, all models benefitted from depth, although the localist model consistently outperformed the GloVe models (Figure 3-B). In terms of predicting human neural dynamics, the localist model peaked at 3 layers, whereas the GloVe based models seemed to benefit more consistently from increased depth (Figure 3-C). However, the localist model still achieved the highest performance overall.

### A modified loss function for human-like lexical competition

Figure 3-D shows lexical competition in the output of the 3 layer localist model during presentation of the target words spoken by the human speaker. For each stimulus, the activation of all words in the lexicon was recorded and sorted by the number of word-initial phonemes shared with the target (cf. Figure 2-H). The target dominates from early on, with little activation of words that share word-initial phonemes. Figure 3-E confirms that the target word dominates lexical activation from early on. Such a pattern may indicate that the model learned idiosyncratic features of individual recordings of words. When models with identical architecture were trained on a different schedule, in which 1000 target tokens were never presented during training (i.e., only different recordings of the same words were used during training), activation of the target was delayed, reflecting the word-initial acoustic ambiguity (Figure 3-F). However, activation of words sharing several onset phonemes with the target remained very low. In contrast, it is thought that humans activate multiple candidates that are compatible with temporarily ambiguous input (Marslen-Wilson, 1987). Activating multiple candidates may be beneficial because it allows evaluating different possibilities, while deferring a final decision until it can be made with reasonable certainty.

The observed pattern in the model output may suggest that the model training schedule prioritized activating a target early on, possibly at the cost of guessing wrong (the cost of guessing is likely lower for the model than it would be for humans because of the smaller vocabulary size). The low level of lexical competition may be a consequence of the binary cross-entropy loss, applied throughout the duration of the word, which primarily rewards activating the correct target as early as possible, and punishes activating competitors. The binary cross-entropy loss effectively imposes a tradeoff between considering multiple words because the loss is minimized when word activations sum to 1. This is sensible for a classification loss where a unique target is required, i.e., the end state of word recognition. However, when information is temporarily ambiguous, it may encourage the model to make premature commitments. In humans, cortical architecture is massively parallel, which may allow considering multiple words matching partial input simultaneously. To develop models that activate cohort competitors, the loss function was modified from the beginning of each word up to 200 ms before the end of the word. During this period, the loss for the target was left unchanged, but the loss for all other words was divided by a constant *c*. Intuitively, the model was still rewarded for activating the right target, but the punishment for activating other words was reduced.

The new loss function led to further improvements in word recognition, with the highest performance at *c*=256 (Figure 3-G). It also dramatically increased the predictive power of deep models for brain activity in the STG (Figure 3-H). As expected, it also led to more realistic activation of phonetic competitors (Figure 3-I), with realistic cohort activation at *c*=256. Models with higher values started indiscriminately activating phonologically unrelated words (*c*=1024, dotted line with peak close to time 0).

### The layers of deep models predict flow of activity from auditory to higher level brain areas

The time course of activity in the different layers of the best performing model reveals an interesting pattern (Figure 4-A). The early layers (L 1-3) track the acoustic input signal, with low activity during periods of silence. The late layers (L 5-6) clearly reflect the target shape, approximating the rectangular on/off wave for periods of word vs. silence. In contrast, in the middle layer 4, the amount of activity does not clearly distinguish between periods of words vs. silence, but contrasting patterns for words vs. silence emerge. This transition was supported by the ability of predicting the amount of activity in higher layers from lower layers (Figure 4-B): Whereas L 2 and 3 were quite predictable given the signal in the layer below, this was not the case for L 4. L 6 was again quite predictable from L 5. This pattern was due to the low frequency modulations: When layer activation was high pass filtered at 4 Hz, each layer was moderately predictable from the layer directly below, and less predictable from layers further away (Figure 4-C).

**Figure 4.**
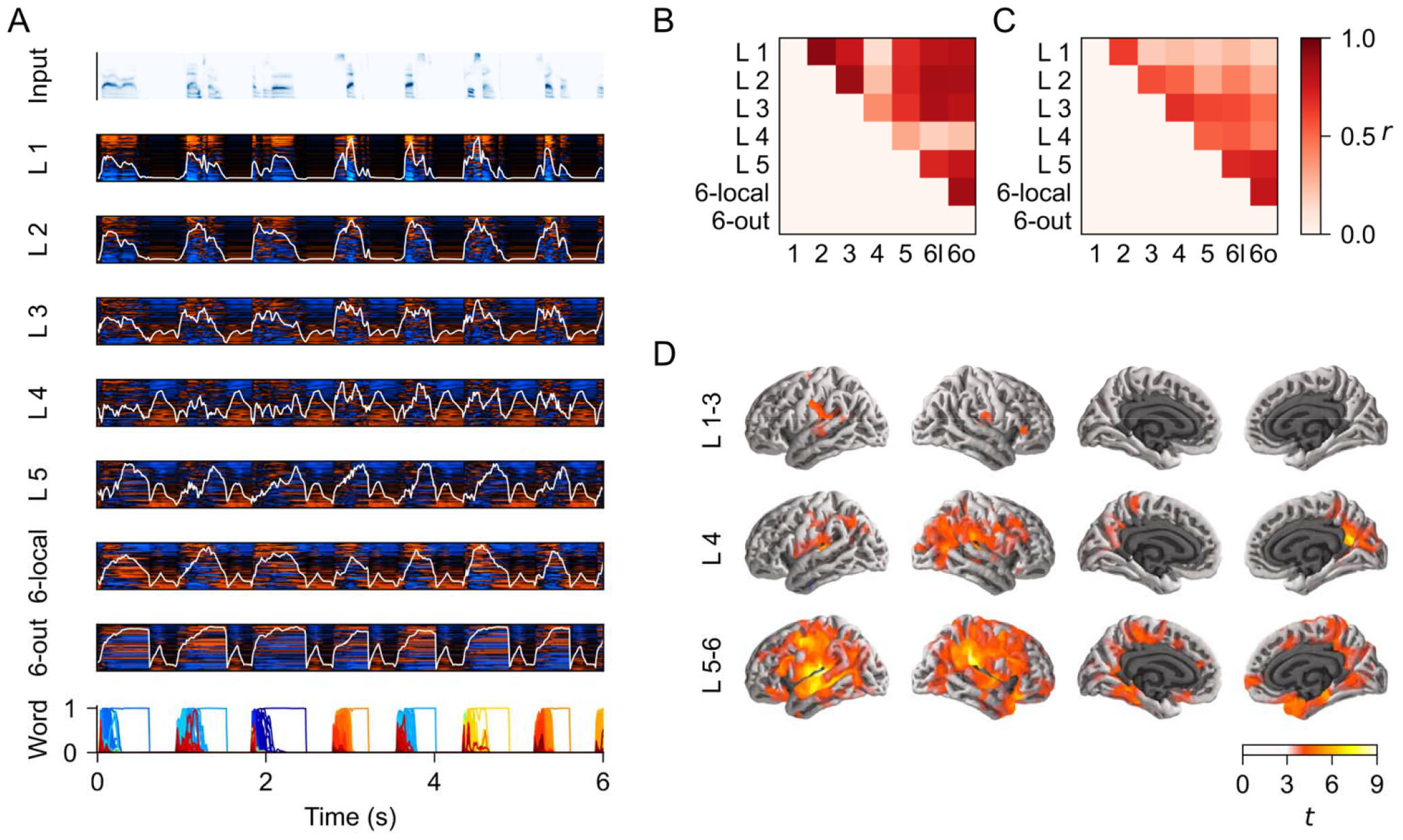
Deep RNN hierarchical activation patterns predict neuroanatomical hierarchy. (A) The first row depicts sample input, and each subsequent row shows activation in a specific layer of the best-performing 6-layer RNN from Figure 3-H. Colors show individual hidden units (sorted by their correlation with the acoustic envelope) and the average magnitude is shown as white line. The last row shows the network output (each line representing activation of one word). ‘6-local’ refers to purely recurrent connections in Layer 6, while ‘6-out’ are hidden units that are connected to the dense output layer. (B) The magnitude signal in the middle layer 4 is more independent than other layers. Each data point reflects the correlation between the signal in one layer (x-axis) predicted with a linear model from the signal in another layer (y-axis). (C) Same as B, but after applying a 4 Hz high-pass filter. (D) The signal in lower layers predicts rain activity close to auditory cortex, whereas the signal in higher layers predicts the signal in higher level brain areas. Each image shows the predictive power of the full model containing all predictors, compared to a model missing the specified predictors, through a mass-univariate *t*-test. Images are masked by significance (whole brain corrected *p* ≤ .05).

A distinction based on hierarchy was also supported by neuroanatomical localization, estimated by comparing the predictive power of the full model to a model in which predictors reflecting activity in specific layers were removed (Figure 4-D). The early layers, which track primarily the acoustic input, predicted brain signals in the close vicinity of the auditory cortex in the superior temporal gyrus; L 4 predicted brain signals in higher level areas including the angular gyrus and medial parietal lobe; and the late layers 5-6 predict brain signals in a wide network of language areas including frontal and temporal areas.

## Discussion

RNNs have been proposed as tools to help us understand the computational demands for human cognitive processes (Elman, 1990), including speech recognition (Magnuson et al., 2020; Yi et al., 2019). Our results demonstrate that this relationship is not just conceptual. RNNs trained to recognize spoken words indeed develop temporal dynamics in their operations that predict human cortical responses to the same words. This suggests that operations learned by those RNNs are in some respects reflective of computations performed for word recognition in the human brain. Critically, altering the details of the task learned by the RNNs alters the predictive power of those RNNs for human brain activity, such that RNNs that are designed to be more consistent with psycholinguistic principles are also more predictive of brain activity. These results suggest that RNNs indeed learn computations that resemble information processing in the human brain.

### Why might RNN activity predict brain activity?

Artificial neural networks, although originally inspired by biology, emulate biological principles very selectively. LSTMs were designed to solve a computational problem, not to imitate how biological networks work (Hochreiter and Schmidhuber, 1997). Nevertheless, hidden unit activity in LSTMs trained to recognize speech can predict the temporal dynamics of neural activity in human listeners. Why might this be? The values in LSTM units store the result of local computations, and make those results available to other units. LSTM units thus transmit information, both forward in time, and to the RNN’s output, or the next layer in hierarchical models. Similarly, human neural activity transmits information within and across regions. Specifically, activity in dendritic trees, the principal generators of the MEG signal, is related to integrating information that is received from other neurons. Thus, the affinity between RNNs and human brain activity may indicate that the two systems develop similar information processing strategies when faced with the same speech signals, and, consequently, that they update internal representations at similar points in time, which leads to similar temporal dynamics in information transmission. This line of reasoning is reinforced by the results showing that when computational demands for RNNs are made more consistent with human cognition, the RNNs also develop more human-like activity patterns. Finally, LSTM internal connectivity may resemble cortical microcircuits (Costa et al., 2017), which may have further promoted similar information processing strategies.

### Neural network activity as a linking hypothesis

A neural linking hypothesis specifies which quantifiable aspects of a cognitive model predict brain activation. In research predicting language-related brain activity from neural network models, typically the values of all hidden units in a specific network layer are used to predict neural population activity (Caucheteux et al., 2023; Li et al., 2023). Here we introduced a novel linking hypothesis, by averaging activation across groups of units. This linking hypothesis is motivated by the physiological basis of M/EEG signals. The electrical activity produced by individual neurons is not strong enough to be measurable above noise levels with M/EEG, but these signals reflect simultaneous activity in populations of spatially aligned cells, i.e., activity in many current vectors that is averaged physiologically (Nunez and Srinivasan, 2006). Even though units in artificial neural networks do not necessarily correspond to biological cells, we hypothesize that our artificial networks developed population dynamics that can be related to biological neural population dynamics. Thus, on this approach, we treat the neural network more like a subject in an electrophysiological experiment, rather than assuming arbitrary access to each individual unit’s activity.

The advantage of the linking hypothesis employed here is that 1) all hidden units of the RNN are incorporated, not just a single layer, and 2) the hidden units within a functional group (layer) are all assigned the same weight for predicting brain activity. The latter is important because in a regression model in which each hidden unit is assigned separate regression weights, the model can selectively use units that resemble brain activity and ignore ones that do not. This is problematic because in such high dimensional models, effects may be driven by a relatively small number of units that happen to resemble human responses by chance. For instance, even untrained deep networks, initialized with random weights, can make surprisingly accurate brain activity predictions (Millet and King, 2021; Schrimpf et al., 2021). This is likely because untrained but structured deep networks provide rich reservoirs of features (Fusi et al., 2016; Jaeger and Haas, 2004), some with response characteristics resembling neural processes *by chance*.

### Lexical activation and competition

During word recognition, human listeners start interpreting partial input early and consider multiple lexical candidates for recognition. For example, upon hearing /bi/, listeners activate lexical representations for *beaker* and *beetle* (Allopenna et al., 1998; Yee and Sedivy, 2006; Zwitserlood, 1989). This is typically conceptualized with a model of the mental lexicon in which each word can be independently activated (Marslen-Wilson, 1987), as is the case in a localist semantic space, where activation of any one word is orthogonal to other words. In a lower-dimensional space, in which words are not orthogonal, the space constrains which words can be activated simultaneously (Gaskell and Marslen-Wilson, 1997; Gaskell and Marslen–Wilson, 1999). In a semantically structured space such as GloVe, activating one word entails activating other words in its semantic neighborhood (e.g., the words closest to *valley* in our GloVe-300 space are *hills, area*, …). However, words that are phonetic onset competitors are not clustered systematically (Figure 2-J). To implement phonetic competition, such a model may point to a location corresponding to an average of the competitors (Gaskell and Marslen–Wilson, 1999). However, this average location may be closer to phonetically unrelated words than to any of the possible targets. For instance, in humans, /væl/ may activate *valley, value, valet* and *valid*, but in our GloVe-300 space the words closest to the arithmetic mean of these four vectors are *hence, example*, and *fact*.

Our results suggest that RNN architectures that allow and facilitate simultaneous activation of phonetic competitors develop the most human-like temporal dynamics. First, models with a trainable target space structured that space according to acoustic-phonetic principles. This may allow them to process the input more efficiently over time by directing their lexical search towards neighborhoods in the target space that contain words with similar onsets. Second, a modified loss function that reduced punishment for activating multiple candidates during early stages of word recognition led to even more human-like temporal dynamics. This may allow for more unconstrained exploration of the lexical space during early stages of word recognition, precluding premature commitments to specific words. In contrast, models that directly mapped sound to the semantic GloVe space made more errors and developed less human-like temporal dynamics. Our results, together with the theoretical considerations above, suggest that a phonetically structured target space is a more appropriate model for human word recognition than a direct mapping to a semantically structures space.

## Conclusions

These results demonstrate that RNNs as models of speech recognition can provide a middle ground between high-level cognitive models and explicit neural implementations. These models learn brain-like computational principles using relatively simple and broadly available computational methods. The success of the LSTM architecture in our investigation adds support for the hypothesis that this architecture implements brain-like computations (Costa et al., 2017).

## Materials and Methods

### Recurrent neural networks

#### Training

The stimuli were based on a set of 1000 target words used in a MEG experiment (Gaston et al., 2023), spoken by a human male speaker for the massive auditory lexical decision (MALD) database (Tucker et al., 2019). To simulate realistic lexical neighborhoods for those target words, additional words were added to the lexicon if they 1) differed from a target word by a single phoneme, 2) had a frequency count of at least 1000 in the SUBTLEX-US corpus (Brysbaert and New, 2009), and 3) were included in the MALD database. This resulted in a total of 2985 words. For each of these words, 15 additional tokens were produced using Apple “say” speakers (Agnes, Allison, Bruce, Junior, Princess, Samantha, Tom, Victoria, Alex, Ava, Fred, Kathy, Ralph, Susan, Vicki). All tokens were transformed to 64 band gammatone spectrograms, roughly corresponding to the frequency resolving power of human hearing (Milekhina et al., 2018) and allowing visually smooth formant movements (see Figure 1). Model training and analysis was performed at 100 Hz temporal resolution.

For RNN training, tokens were divided into training and test sets. For each of the 16 speakers, 1/16^th^ of the tokens were randomly assigned to the test set, such that each word was trained with 15 tokens from 15 different speakers, and with one token/speaker held out for testing. To minimize item variability when predicting MEG data from RNNs, the randomization was constrained to always include the MALD tokens presented in the MEG experiment in the training set.

Training input consisted of a continuous stream of words and silence. The following rules were used to mimic the continuity in the input due to speakers commonly producing sentences rather than isolated words: 1) select a speaker; then, produce two phrases according to: 2) select a token from the speaker; 3) with a 50% chance, go back to (2); 4) insert a silence of 200-500 ms. Return to (1).

All RNN models were trained with TensorFlow 2.8 (TensorFlow Developers, 2023). Inputs were created by concatenating gammatone spectrograms and silence into 10 s long segments, but implemented a fully continuous time axis: for example, a word might start at the end of one segment and continue in the beginning of the next segment, and LSTM states carried over across segments. Gaussian noise was added to all stimuli at 20 dB signal-to-noise ratio (determined separately for each frequency band). Models were trained in parallel with a batch size of 32. Each training epoch consisted of 25 10 s segments. Models were trained to minimize binary cross-entropy (localist models) or mean squared error (all other models) across *all* time points. Training loss was evaluated after each epoch. When the loss increased over a previous minimum for 20 epochs, model weights were restored to that previous minimum. When the loss failed to improve for 200 samples, model training was terminated. In a few instances, this led to models that did not converge (>99% word error rate). In those cases, models were retrained with slightly modified parameters: epoch lengths were increased to 50 10 s segments, and model training was stopped after the loss failed to improve for 250 epochs. This led to successful training in all cases.

#### Evaluation

For model evaluation, the trained models were presented with a continuous sequence of all tokens without any intervening silence, and without added noise. Training set tokens were presented before the test set tokens; otherwise, the order was completely randomized (without replacement). To evaluate model performance, each model’s output was transformed into lexical activation (i.e., a vector assigning a value to each word, where the largest value can be interpreted as the word that the model is currently perceiving). Localist models directly output a vector of activation for each possible word. For semantic vector space models, word activation was defined as the negative distance of the output from each word in the lexicon (i.e., the model’s percept was defined as the word in the lexicon with the smallest distance from the output). For SRV models, the output was transformed into lexical activation by assigning to each word an activation value corresponding to the minimum across the 10 output elements corresponding to that word. Word error rate was then determined based on the average word activation (as defined above) during the last 100 ms (i.e., 10 samples) of the target word. The word was recognized correctly only if the most highly activated word corresponded to the actual target. Word error rate is generally reported for the test set only, encompassing 2985 tokens (one instance of each word).

### MEG dataset and analysis

MEG data were drawn from a public dataset (Gaston et al., 2022). Preprocessing and source localization are summarized here, and are described in more detail in a previous publication (Gaston et al., 2023). The analysis included data from 18 right-handed, native speakers of English. Participants listened to 1000 words taken from the MALD database (Tucker et al., 2019), with an interstimulus interval of 276 ms. Occasionally, a visual probe word appeared on the screen and participants judged whether the word was semantically related to the word they had heard most recently, and all participants included in the analysis answered at least 69% of probes correctly.

#### Preprocessing

Extraneous artifacts were automatically removed using temporal signal space separation (Taulu and Simola, 2006). Data were then band-pass filtered between 1 and 40 Hz, and biological artifacts were removed using extended infomax independent component analysis (Lee et al., 1999). Data were then low-pass filtered at 20 Hz and downsampled to 50 Hz.

A source space for MEG analysis was defined on the white matter surface of the FSAverage template brain (Fischl, 2012). This source space was uniformly scaled for each participant to approximate their head size. MEG data for the whole experiment were projected to virtual current sources, oriented perpendicularly to the cortical surface, using regularized minimum *ℓ*2 norm estimates (λ= 1/6).

#### Auditory baseline model

A model of auditory processing was generated based on gammatone spectrograms as in previous experiments (Brodbeck et al., 2020; Gaston et al., 2023). First, high resolution gammatone spectrograms were generated for all word stimuli with 256 frequency bands in equivalent rectangular bandwidth (ERB) space between 20 and 5000 Hz. These spectrograms were resampled to 1000 Hz log-transformed to better approximate response characteristics of the auditory system (Brodbeck et al., 2023). Onset spectrograms were generated by applying a neurally inspired edge detector algorithm (Brodbeck et al., 2020; Fishbach et al., 2001). Spectrograms and onset spectrograms were downsampled to 8 frequency bins, equally spaced in ERB space, for mTRF model estimation.

In addition to auditory predictors, the baseline model also included a predictor with a single impulse at each word onset, to account for any generic onset processing not captured by the acoustic predictors.

#### RNN linking hypothesis

The basic linking hypothesis we employed assumes that non-zero values in LSTM hidden units correspond to neural activity. To implement this, a predictor variable was defined as the sum of the absolute values of all hidden units in a layer, at each time point. A variation of this linking hypothesis assumes that biological neural activity may correspond to updates in RNN hidden units, rather than raw values (Rabovsky et al., 2018). To implement this, a second predictor variable was computed by taking the derivative in time of the absolute values of each unit, applying half-wave rectification, and then summing across units. Half-wave rectification, i.e. clipping negative derivative values to 0, assumes that hidden units moving away from zero represent new information, whereas returning to zero constitutes a return to baseline (compare hidden unit activities in Figure 1). Pretests suggested that both predictor types contributed independently to predictive performance of the models. In the reported tests we always include both predictor types, i.e., each LSTM layer was always represented by two predictor variables, reflecting magnitude and change of hidden unit activity.

#### Multivariate temporal response function (mTRF) analysis

Source localized MEG responses were analyzed using multivariate temporal response functions (mTRFs) with Eelbrain (Brodbeck et al., 2023). Separate mTRF models were estimated for each set of predictors and for each participant. Each dataset (MEG data and time-aligned predictors for one subject) was analyzed as one continuous recording. First, the data were divided in 4 contiguous segments for 4-fold cross-validation. For each of the 4 test segments, the three remaining segments were used to estimate 3 different mTRF models, with each segment serving as validation segment once and as training segment in the other two runs.

mTRFs were estimated with a mass-univariate approach, independently estimating mTRFs for each virtual current source in the neural source space. In general, an mTRF model estimates the MEG signal time series at a virtual current source, *ŷ_t_*, as a linear convolution of *N* predictor variable time series, *x_i,t_*, with the mTRF *h_i,τ_*, where τ indexes one of T time delays between predictor variable and brain response:

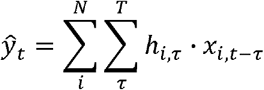

In the current analysis, *τ* always ranged from −100 to 1000 ms. Within each run, a coordinate descent algorithm was used to minimize the ℓ2 error: First, the training data segments were used to find the mTRF element *h_i,τ_* which, when changed by a fixed step Δ, led to the smallest error in the training data. Then, the validation segment was used to test whether the new mTRF also led to an improvement in the validation data. If new change led to an increase in validation error, then *h_i,τ_* was reset to its previous value, and the mTRF components for predictor *i* were frozen. This proceeded until the whole mTRF was frozen.

The 3 mTRFs for each test segment were averaged and used to predict the MEG responses in the test segment. The test segments were then concatenated to compute the fit metric (% of the variance explained) for the whole dataset.

### Statistics

Statistics are implemented in Eelbrain (Brodbeck et al., 2023). For ROI tests, an ROI was defined based on the combination of the anatomical labels for the superior temporal gyrus and the transverse temporal gyrus in each hemisphere (Desikan et al., 2006). The % variance explained was averaged in this ROI and analyzed with standard univariate tests (related measures *t*-tests and ANOVA). For mass-univariate *t*-tests (Figure 4), family-wise error correction was implemented with threshold-free cluster enhancement (Smith and Nichols, 2009), with a null distribution estimated from 10,000 random permutations of the condition labels.

## Acknowledgements

We thank Kevin Brown, Emily Myers and Phoebe Gaston for discussions on related topics. This work was supported by National Science Foundation grants BCS-2043903 and IIS-2207770 to JSM and CB. JSM’s effort was also supported in part by the Basque Government through the BERC 2022-2025 program and by the Spanish State Research Agency through BCBL Severo Ochoa excellence accreditation CEX2020-001010-S and through project PID2020-119131GB-I00 (BLIS).

